# Specificities and commonalities of carbapenemase producing *Escherichia coli* isolated in France from 2012 to 2015

**DOI:** 10.1101/2021.10.19.464995

**Authors:** Rafael Patiño-Navarrete, Isabelle Rosinski-Chupin, Nicolas Cabanel, Pengdbamba Dieudonné Zongo, Mélanie Héry, Saoussen Oueslati, Delphine Girlich, Laurent Dortet, Rémy A Bonnin, Thierry Naas, Philippe Glaser

**Author notes:** Corresponding author: Philippe GLASER, Institut Pasteur, 28 Rue du Dr Roux, 75724 Paris Cedex 15. Contributed equally (Thierry Naas, Philippe Glaser). Contributed equally (Rafael Patiño-Navarrete, Isabelle Rosinski-Chupin).

## Abstract

Carbapenemase-producing *Escherichia coli* (CP-*Ec*) represent a major public health threat with a risk of dissemination in the community as it has occurred for lineages producing extended spectrum ß-lactamases. To characterize the extend of CP-*Ec* spread in France, isolates from screening and infection samples received at the French National Reference Centre laboratory (F-NRC) for carbapenemase-producing Enterobacterales were investigated. Six hundred and ninety one CP-*Ec* isolates collected between 2012 and 2015 and 22 before were fully sequenced. Analysis of their genome sequences revealed some disseminating multidrug resistant (MDR) lineages frequently acquiring diverse carbapenemase genes mainly belonging to clonal complex (CC) 23 (ST 410) and CC10 (ST10, ST167) and sporadic isolates including rare ST131 isolates (n=17). However, the most represented ST was ST38 (n=92) with four disseminated lineages carrying *bla*_OXA-48-like_ genes inserted in the chromosome. Globally, the most frequent carbapenemase gene (n=457) was *bla*_OXA-48_. It was also less frequently associated with MDR isolates being the only resistance gene in 119 isolates. Thus, outside the ST38 clades, its acquisition was frequently sporadic with no sign of dissemination, reflecting the circulation of the IncL plasmid pOXA-48 in France and its high frequency of conjugation. In contrast *bla*_OXA-181_ or *bla*_NDM_ genes were often associated with the evolution of MDR *E. coli* lineages characterized by mutations in *ftsI* and *ompC*.

**IMPORTANCE:** Carbapenemase-producing *Escherichia coli* (CP-*Ec*) might be difficult to detect, as minimal inhibitory concentrations can be very low. However, their absolute number and their proportion among carbapenem-resistant *Enterobacterales* have been increasing, as reported by WHO and national surveillance programs. This suggests a still largely uncharacterized community spread of these isolates. Here we have characterized the diversity and evolution of CP-*Ec* isolated in France before 2016. We show that carbapenemase genes are associated with a wide variety of *E. coli* genomic backgrounds and a small number of dominant phylogenetic lineages. In a significant proportion of CP-*Ec*, the most frequent carbapenemase gene *bla*_OXA-48_, was detected in isolates lacking any other resistance gene, reflecting the dissemination of pOXA-48 plasmids, likely in the absence of any antibiotic pressure. In contrast carbapenemase gene transfer may also occur in multi-drug resistant *E. coli*, ultimately giving rise to at-risk lineages encoding carbapenemases with a high potential of dissemination.

## INTRODUCTION

*Escherichia coli* is one of the first causes of diverse bacterial infections in the community and in the hospital. In particular, it is the most frequent cause of urinary tract infection (UTI) and 50-60% of women will suffer at least once a UTI during her life (1). Therefore, multidrug resistance (MDR) in *E. coli* is a major public health issue making *E. coli* infections more difficult to treat. In addition, as carbapenemase-producing *Enterobacterales* (CPE) are increasingly detected (2, 3) and *E. coli* is a ubiquitous member of the human gut microbiome, carbapenemase-producing *E. coli* (CP-*Ec*) are also becoming a major actor for the global dissemination of carbapenemase genes (4).

The emergence and spread of carbapenem-resistant Gram-negative bacteria is mainly linked to the widespread dissemination through horizontal gene transfer (HGT) of mobile genetic elements (MGEs) encoding carbapenemases. These carbapenemases belong to Ambler class A (mainly KPC-type), class B (metallo-β-lactamases IMP-, VIM- and NDM-types) or class D (OXA-48-like enzymes) of beta-lactamases (5). The global epidemiology of extended spectrum ß-lactamases (ESBL) producing *E. coli* has been characterized through multiple studies, revealing in Western countries the major contribution of the sequence type (ST)131 lineage in the high prevalence of *bla*_CTX-M_ family ESBL genes (6). Much less is known with respect to CP-*Ec*.

Since in 2012, the French National Reference Centre Laboratory for carbapenemase-producing *Enterobacterales* (F-NRC) has experienced a steady increase in the number of CP-*Ec* isolates received each year (2). A multi locus sequence typing (MLST) analysis of isolates received in 2012 and 2013 revealed a broad diversity of STs, as the 140 analyzed isolates belonged to 50 different STs. However, a few STs were over-represented (7), such as the ST38 (24 isolates) and the ST410 (10 isolates). In that study, only one isolate belonged to ST131, contrasting with the situation in the UK where ST131 isolates represented 10% of the CP-*Ec* isolates received between 2014 and 2016 by Public Health England (8). On the other hand, a genome based survey of CPE in the Netherlands between 2014 and 2018, revealed that the 264 analyzed *E. coli* isolates belonged to 87 different STs, with three dominant lineages, ST38 (n=46), ST167 (n=22) and ST405 (n=16) representing 32 % of the isolates (9). F-NRC isolates also showed a predominance of isolates producing OXA-48-like carbapenemases followed by NDM-producing isolates and suggested clonally related isolates among ST38 OXA-48-producing isolates and ST410 OXA-181-producing isolates, respectively (7).

Whole-genome sequencing (WGS) significantly increases our ability to infer phylogenetic relationships between isolates. By sequencing 50 ST410 CP-*Ec* isolates received by the F-NRC between 2011 and 2015 (10) we showed that 36 (72%) belonged to a newly described ST410 lineage producing OXA-181 (11). We showed that this clade is characterized by mutations in the two porin genes *ompC* and *ompF* leading to a decreased outer membrane permeability to certain ß-lactams and by a four-codon duplication (YRIN) in the *ftsI* gene encoding the penicillin binding protein 3 (PBP3) leading to a decreased susceptibility to ß-lactams targeting this PBP (10). After a thorough analysis of CP-*Ec* genome sequences from public databases for mutations in these three genes, we proposed that CP-*Ec* followed three different evolutionary trajectories. In some lineages which are enriched in CP-*Ec* isolates and have disseminated globally, acquisition of carbapenem resistance genes might have been facilitated by the mutations in porin genes and in *ftsI*. In ST38, the genetic background and in particular a specific *ompC* allele with reduced permeability to some ß-lactams may have similarly facilitated the acquisition of carbapenemase genes. In contrast, other CP-*Ec* isolates including from ST131 might have occurred sporadically following the acquisition of plasmids encoding carbapenemase and with no clue of dissemination (10).

Here we thoroughly characterized the diversity of CP*-Ec* circulating in France by sequencing the genome of almost all isolates received by the F-NRC from its creation in 2012 until 2015 (Table S1). By combining whole genome phylogeny with the addition of *E. coli* genome sequences publicly available (Table S2), and tracking mutations in candidate genes, we show that three different situations are encountered. The high transmission potency of the IncL pOXA-48 plasmids has led to a high frequency of OXA-48-producing isolates, often characterized by susceptibility to most non-ß-lactam antibiotics. On the other hand, an increasing number of CP-*Ec* lineages, mainly producing OXA-181 and NDM carbapenemases, are observed in France as in other European countries. These lineages are multi-drug (MDR) or extensively drug resistant (XRD) lineages, they show several mutations in quinolone resistance determining regions (QRDR) and are often mutated in *ftsI* and in porin genes. Finally, the rapid dissemination of four OXA-48/OXA-244 ST38 lineages might have been favored by the chromosomal integration of the carbapenemase gene.

## RESULTS

### CP-*Ec* isolates collected until 2015 by the French National Reference Center

Isolates analyzed in this work were sent to the F-NRC laboratory on a voluntary basis from private and public clinical laboratories from different parts of France mainly between the years 2012 to 2015. During this period, we encountered a regular increase in the number of isolates received each year (Fig 1A) corresponding to an increasing number of isolated strains and an increasing frequency of isolates sent to the F-NRC. 713 CP-*Ec* isolates, including 22 collected from 2001 to 2011 were submitted to WGS. The 691 sequenced isolates of the 2012-2015 period represented 87.5 % of the 790 CP-*Ec* isolates received by the F-NRC during this period.

**Figure 1.**
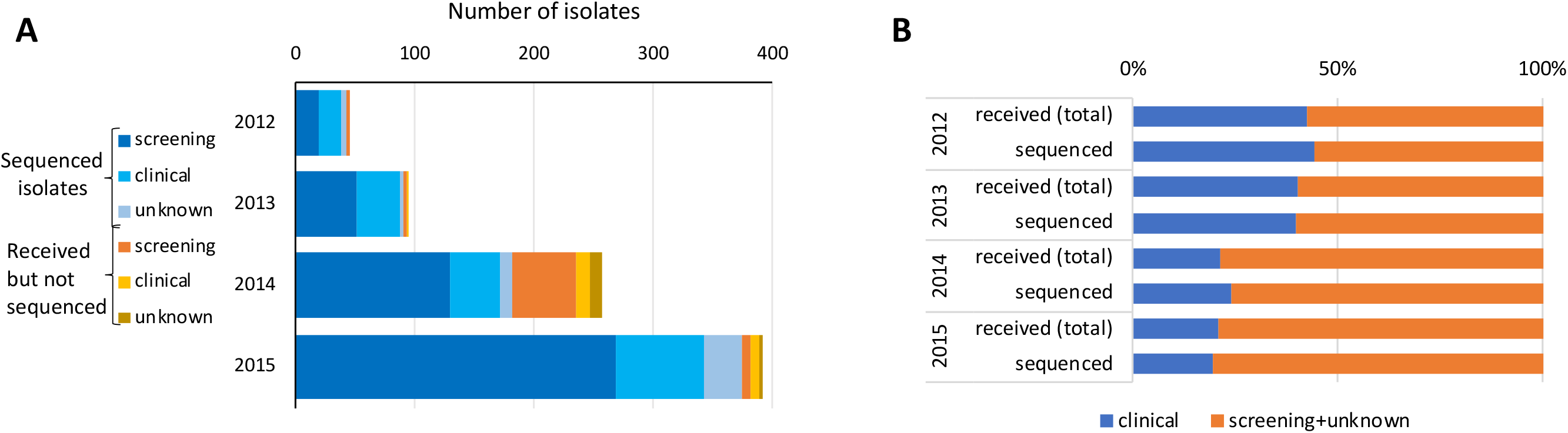
Origin of the isolates received by the F-NRC during the 2012-2015 period. **A**. Absolute number of isolates per year from infection, screening or unknown origins, differentiating between isolates that have or not been sequenced. **B**. Proportions of isolates from infection or from screening or unknown origins showing that isolates from infection origin represent the same proportions among total isolates that have been received by the F-NRC and isolates that have sequenced.

The majority of the sequenced isolates (66.5%, 474/713) were from rectal swab screening of patients suspected of carrying CPE (patients repatriated from an hospital abroad, patients that have visited a foreign country within the last six months, contact patients of a former carrier, or a previously known carrier), 24% (172/713) were considered to be responsible for infection (isolated from clinical samples), and the source was unknown for 67 isolates (Fig. 1A, Table S1). During the four-year period, The number of infection-related isolates was found to increase more slowly than the number of screening or of unknown source isolates. A nearly two-fold decrease in the proportion of clinical versus total isolates was observed between 2013 and 2014 (Fig. 1B). Among the 172 isolates associated with disease, 122 were isolated from urine (71%), 15 from blood samples, eight from deep site samples, eight from wound samples, nine from the respiratory tract, three from vaginal samples, three from pus, three from bile, and one from the skin of a newborn. All these isolates were previously identified as carbapenemase producers by PCR.

### Diversity of CP-Ec isolates as assessed by whole genome sequencing

The genome sequences of the 713 CP-*Ec* isolates were first analyzed following *de novo* assembly. For each isolate, we determined its MLST type (Achtman scheme), its phylogroup (ClermonTyping) and its antibiotic resistance genes (ARG) content. We also searched for mutations in the QRDRs of *gyrA*, *parC* and *parE,* and for mutations in *ftsI*, *ompC* and *ompF* we previously identified as associated with CP-*Ec* disseminated lineages (10) (table S1). F-NRC CP-*Ec* were assigned to 168 different STs including six new allelic combinations. ST38 was the most prevalent ST (n=92, 12.9 %), followed by ST10 (n=67, 9.4 %), ST410 (n=64, 9 %), and ST167 (n=34, 4.8 %). Ten additional STs were represented by at least 10 isolates (Fig. S1A, table S1) and 154 STs with less than ten isolates including 102 STs with a single isolate.

We next performed a core genome phylogeny following read mapping using strain MG1655 (NC_000913.3) as reference genome sequence. The phylogenetic tree, estimated from 372,238 core SNPs was consistent with the results of phylogroup determination by using *in silico* ClermonTyping (12) except for a few isolates (Fig. 2). In agreement with the MLST-based analysis, this tree showed a broad diversity of CP-*Ec* isolates belonging to the eight phylogroups and three dominant clades corresponding to CC10 (phylogroup A; including ST10, ST167 and ST617), CC23 (phylogroup C; including ST410, ST88 and ST90) and ST38 (phylogroup D) with 161, 97, and 98 isolates respectively, accounting for 49.9 % of the F-NRC CP-*Ec* isolates analyzed (Fig. 2). Phylogroup B2 isolates represented 11 % of the total isolates (n=80) and only 17 CP-ST131 isoaltes were identified. Fluctuations in the proportions of the main STs could be observed during the four years of the analysis but no clear tendency could be identified (Fig. S1A).

**Figure 2:**
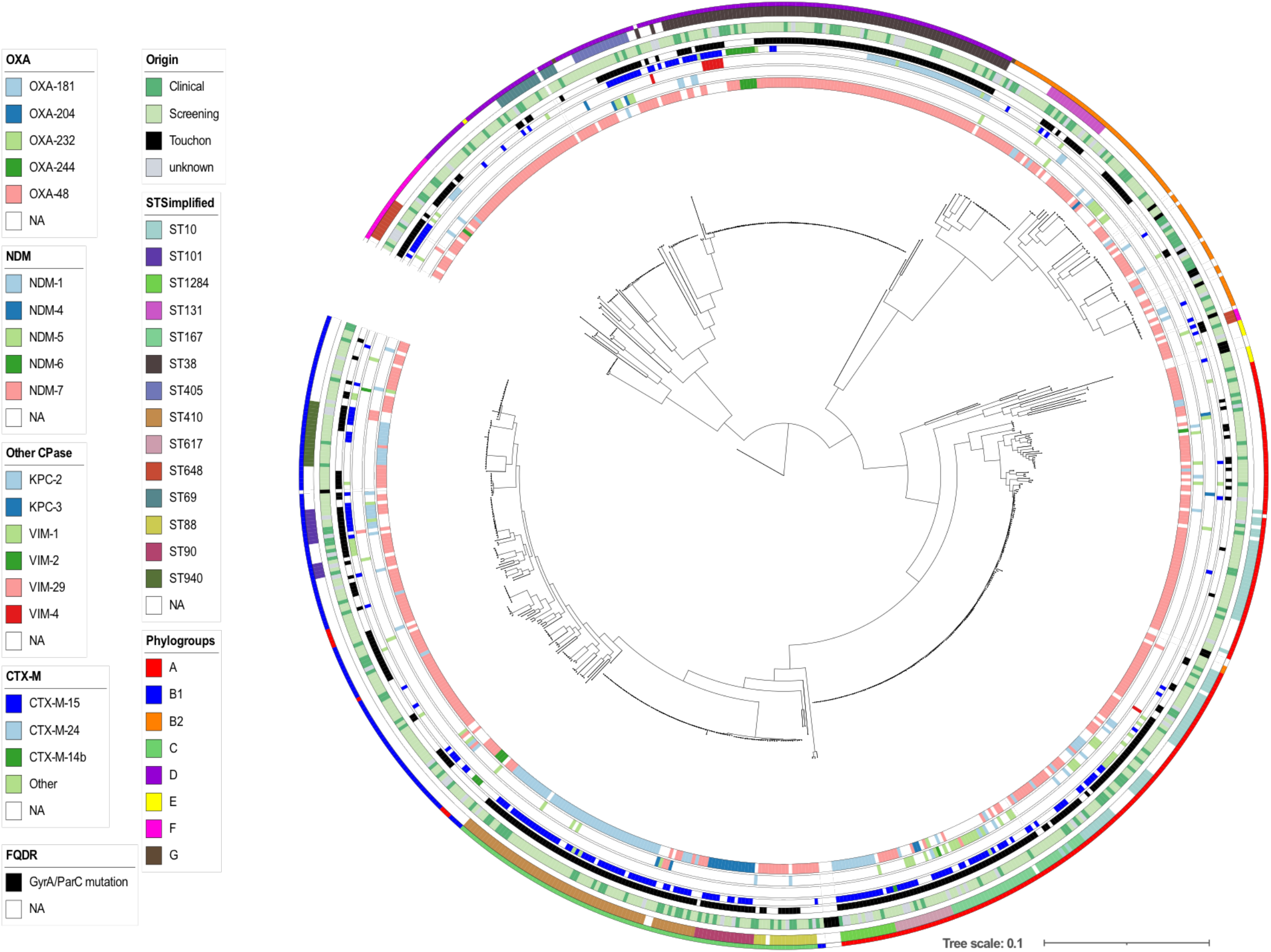
Core genome phylogeny of CP-*Ec* isolates received by the F-NRC. ML phylogeny of the 713 CP-*Ec* isolates was built with RAxML (45) from 372,238 core SNPs after sequence alignment on MG1655-K12 (NC_000913.3) selected as the best reference. The genome sequence of *Escherichia fergunsonii* strain ATCC 35469 (NC_011740.1) was used as an outgroup for the phylogenetic analysis. Genomes from Touchon et al. (40) were also incorporated into the analysis. Genomic features are indicated as in the figure key (left) from the inside to the outside circles: carbapenemases of the OXA, NDM and other types, CTX-M ESBL, mutations in *gyrA* and *parC* QRDR region (FQ resistance), origin, main ST, phylogroups. Reference genome from Touchon *et al*. are indicated as Touchon.

Infection-related and screening isolates were intermixed throughout the phylogeny (Fig. 2). However, an enrichment in infection-related isolates was observed in phylogroup C (Pearson’s Chi-squared test, p<0.02, ddl1) and phylogroup B2 (Pearson’s Chi-squared test p<0.0005, ddl1) (Fig. S1B). In phylogroup B2, 5 out of 9 ST127 isolates, 5 out of 17 ST131 isolates and 7 out of 8 ST636 were responsible for UTIs (Table S1).

### Diversity of antibiotic resistance genes carried by CP-Ec isolates

The number of acquired ARGs was found to vary between 1 and 26 among the 713 CP-*Ec* isolates. The median was higher in phylogroup C isolates (m=16) and lower in isolates from phylogroup B2 (m=3) compared to other phylogroups (Medians: A: 9; B1: 9; D: 11) (Fig. 3A). An ESBL of the CTX-M family was present in 40.7% (N=290) of the isolates, with a predominance of *bla*CTX-M-15 gene (N=205) (Table S1). Mutations in g*yrA*, *parC* and/or *parE* potentially leading to fluoroquinolone (FQ) resistance occurred in 425 CP-*Ec* isolates (59.6%), with mutations in *gyrA*, *parC* and *parE* identified in 412 isolates, 309 and 261 isolates respectively (Table S1). Up to five mutations in QRDRs were identified in 19 isolates and 250 isolates had four mutations in QRDR, suggesting they have been submitted to a long-term evolution under antibiotic pressure including FQ (Table S1). Globally a higher number of mutations in QRDR was associated with a higher number of resistance genes (Fig. 3B), furthering the link between the number of QRDR mutations and a likely evolution under antibiotic selective pressure for CP-*Ec* isolates.

**Figure 3.**
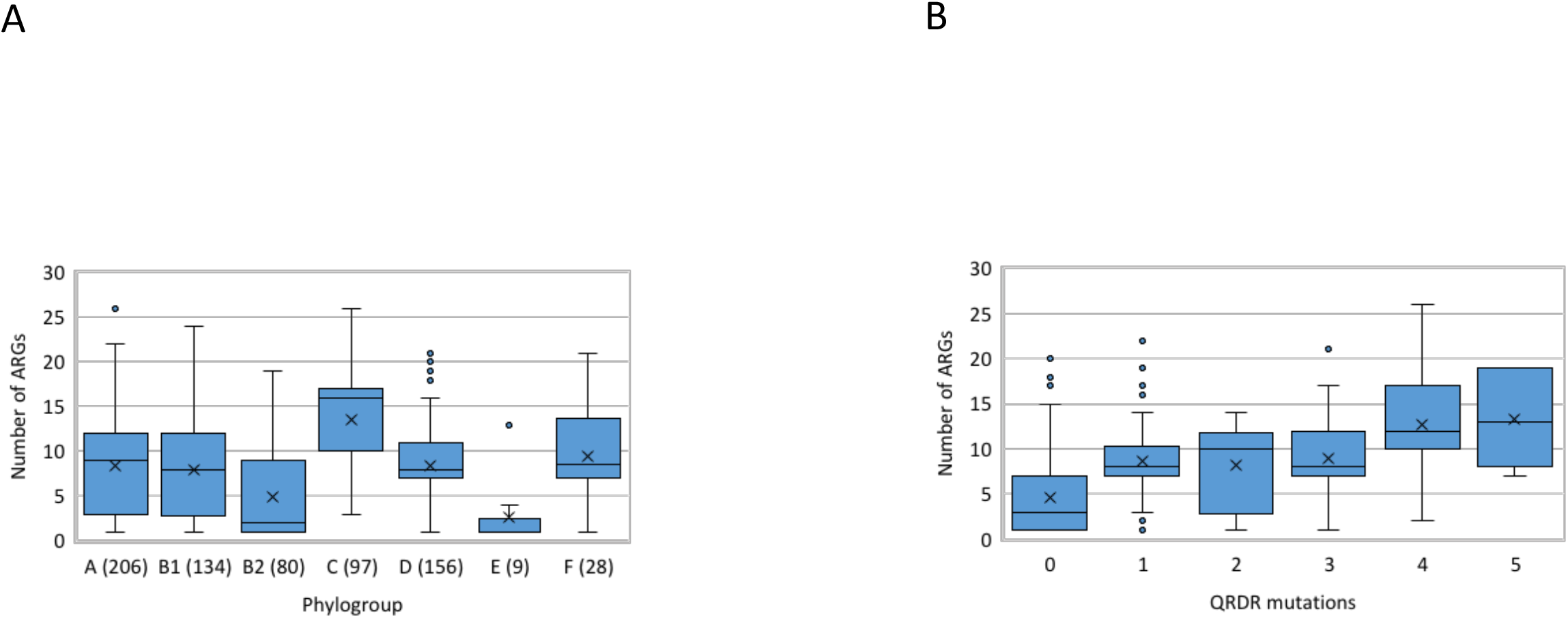
Resistance gene content of the F-NRC CP-*Ec* isolates. **A**. Box plot representation of the ARGs content according to the phylogroup. The number of isolates belonging to each phylogroup is indicated between brackets. The limits of the box indicate the lower and upper quartile. Outliers are indicated by points above the maximal value. **B**: Relationships between the ARGs number and the number of mutations in QRDR. In the box plot representations, the median is indicated by an horizontal bar and the mean by a cross.

Four isolates from 2015, two ST648 collected in a same hospital at one-week interval, one ST216 and one ST744, encoded a *mcr-9* gene conferring resistance to colistin (Table S1). Although not associated with infection, the four isolates were MDR, carrying 13 to 20 acquired ARGs in addition to *mcr-9* and carbapenemase genes (*bla*_VIM-1_ for three of them, *bla*_NDM-1_ and *bla*_OXA-48_ for one). Determination of the MIC for colistin for these four isolates were at the resistance breakpoint (MIC=2 mg/l).

### Diversity of carbapenemase genes

Carbapenemase genes identified by WGS (Table S1) were in agreement with molecular data collected by the F-NRC laboratory. The *bla*_OXA-48_ and *bla*OXA-181 genes were the most frequent carbapenemase genes detected in 464 (65%) and 101 (14.1%) *E. coli* isolates respectively. Twenty-nine other isolates carried minor *bla*_OXA-48_-like genes (Table S1). *bla*_NDM_ family genes were detected in 14.9% (106/713) of the CP-*Ec* isolates (*bla*_NDM_-5 in 49 isolates and *bla*_NDM_-1 in 41). Three additional *bla*_NDM_ alleles were identified: *bla*_NDM_-7 (n=10) including a variant coding for a NDM7-like carbapenemase with a S24G mutation, *bla*_NDM_-4 (n=5), and *bla*_NDM_-6 (n=1). Four different *bla*VIM alleles were detected in 18 isolates: *bla*VIM-1 (n=8), *bla*VIM-4 (n=8), *bla*VIM-2 (n=1), and *bla*VIM-29 (n=1). Only five *E. coli* isolates expressed *bla*KPC-_2_ (n=2) or *bla*_KPC-3_ (n=3) alleles. In twelve isolates, two different carbapenemase genes were found; *bla*_OXA-48_ / *bla*_NDM-1_ was the most frequent co-occurrence (n=5) (Table S1).

The analysis of the number of additional ARGs and of mutations in QRDR regions showed that, compared with other carbapenemase genes the presence of *bla*_OXA-48_ was frequently associated with less resistant isolates (Fig. 4A, 4B). In particular it was the only ARG in 117 isolates among the 457 (24.5%) encoding this carbapenemase, including 23 infection-related isolates. Among those isolates, only five showed mutations in QRDR. Conversely, only two *bla*_NDM-1_ gene carrying isolates among 42 and one *bla*_NDM-5_ gene carrying isolates among 49 carried no other ARG.

**Figure 4.**
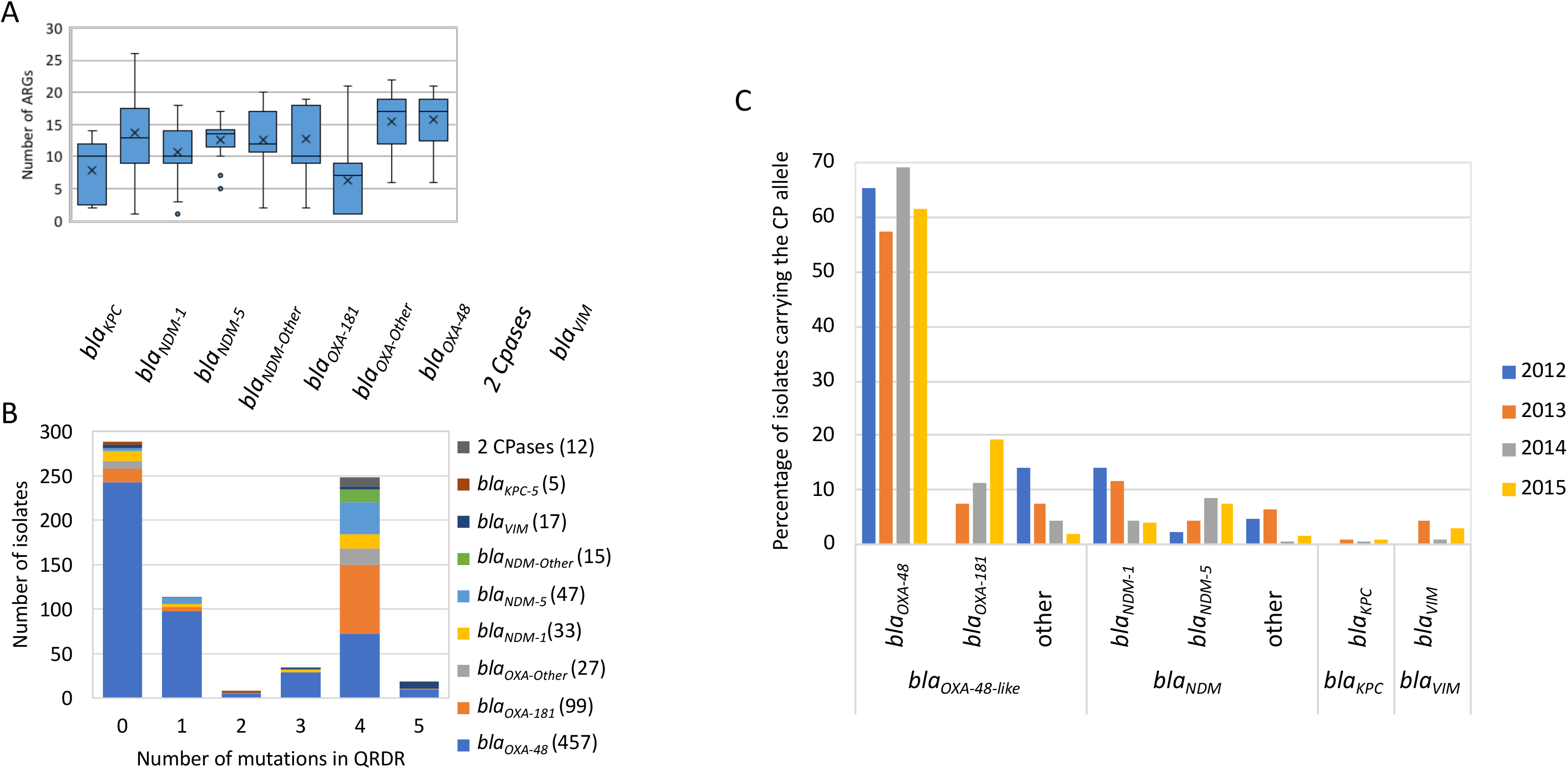
Carbapenemase gene content of the F-NRC CP-*Ec* isolates. **A**. Relationship between the carbapenemase allele and the ARGs content. In the box plot representation, the median is indicated by an horizontal bar and the mean by a cross. The limits of the box indicate the lower and upper quartile. Outliers are indicated by points above the maximal value. **B**. Number of isolates carrying a specific carbapenemase allele according to the number of QRDR mutations in their genomes. **C**. Per-year evolution of the percentage of isolates carrying the different carbapenemase alleles in the 2012-2015 period. Year of isolation are indicated as in the figure key (right). Given their small number (n=22), strains isolated before 2012 are not indicated.

A temporal analysis of the proportion of isolates displaying *bla*_OXA-48_ did not reveal significant variations (Pearson’s Chi-squared test, p=0.05, ddl= 3) during the 2012-2015 period (Fig. 4C). In contrast, the frequency of *bla*_OXA-181_ relatively to other alleles was found to significantly increase from 0 % in 2012 to 18.9% in 2015 (Pearson’s Chi-squared test, p < 0.0002 ddl=3) (Fig. 4C).

We observed some association between the carbapenemase gene and the ST. Among the 22 STs with at least seven F-NRC CP-*Ec* isolates, fourteen predominantly displayed *bla*_OXA-48_ (Fig. 5A); *bla*_OXA-48_ was also predominant in STs represented by six or less isolates. In three lineages, ST410, ST940, and ST1284, *bla*_OXA-181_ was the predominant allele. These three STs grouped 74% of the *bla*_OXA-181_ gene carrying *E. coli* isolates from the F-NRC, and ST410 accounted for 50.5% of them. Finally, *bla*_OXA-204_ was the most prevalent allele in ST90 and *bla*_NDM-5_ the predominant allele in ST636.

**Figure 5.**
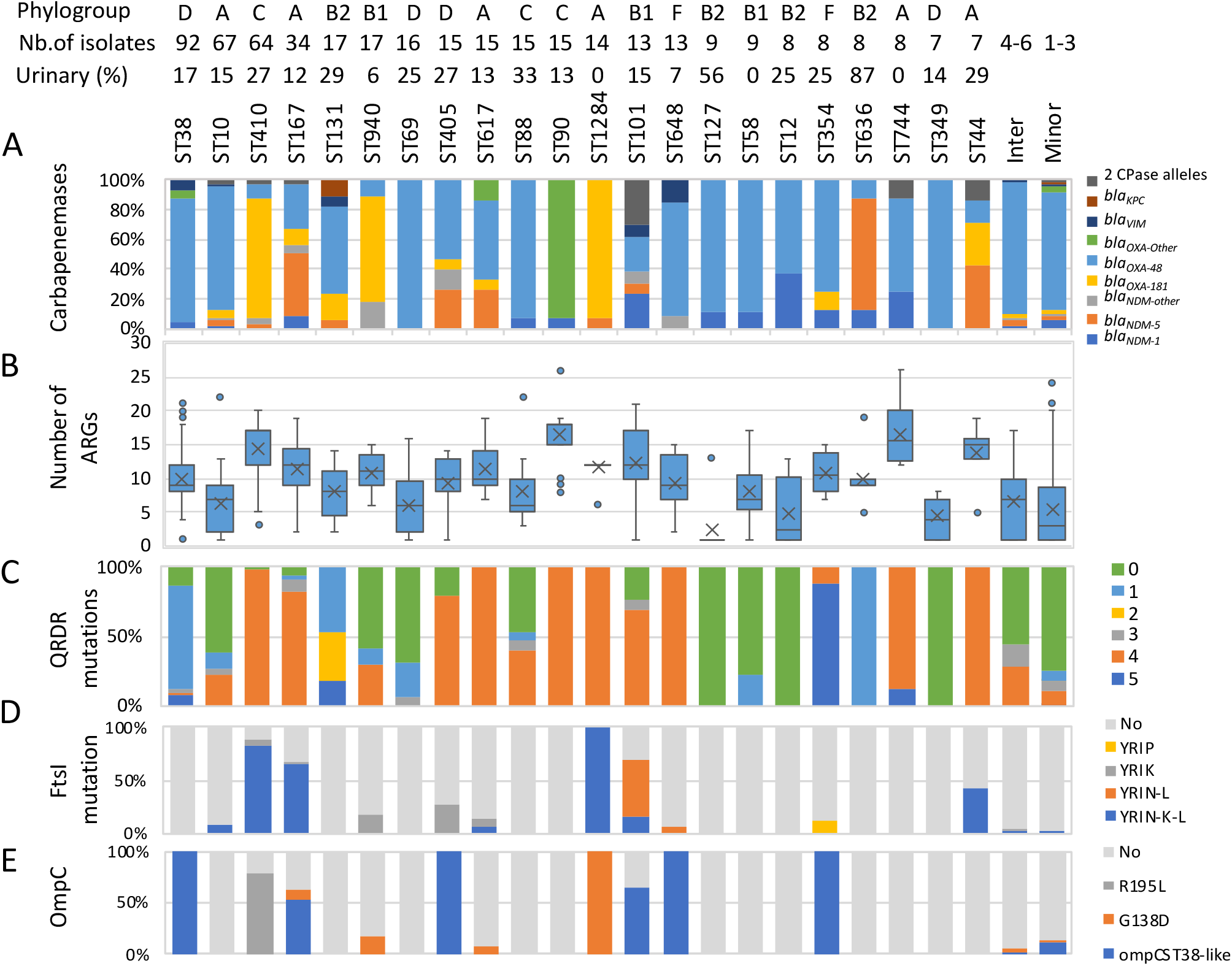
Features of the main ST associated with CP-*Ec* isolates in France. **A**. Proportion of the isolates according to the carbapenemase allele in each ST; **B**. Box plot representation of the ARGs content according to the ST; **C.** Proportion of isolates according to the number of mutations in QRDR; **D**: Proportion of isolates with a specific PBP3 allele. PBP3 characterized by a YRIP, YRIK or YRIN insertion between positions 333 and 334, the YRIN insertion is generally associated with a A413V mutation (YRIN-L) and less frequently with a E349K mutation (YRIN-K-L) (10); **E.** Proportion of isolates carrying specific mutations in *ompC:* R195L, G138D or acquisition of a ST38-like *ompC* allele by recombination (10). For each ST, the phylogroup, the total number of isolates and the number of isolates associated with urinary tract infections are indicated in the upper panel. Intermediate (Inter) and minor ST indicate ST represented by 4-6 and 1-3 isolates, respectively.

### Characteristics of the STs preferentially associated with F-NRC CP-Ec isolates

Eight among the 14 STs with more than 10 CP-*Ec* isolates (ST410, ST167, ST940, ST405, ST617, ST90, ST1284, ST101) were characterized by a larger number of ARGs (median ≥ 10) and a larger number of mutations in QRDR as compared to ST38, ST10, ST131, ST69, ST88 and ST represented by less than seven isolates (Fig. 5 B, C). ST648 isolates have an intermediate position, being highly mutated in QRDR but less rich in ARGs. Analysis of CP-*Ec* isolates from the F-NRC for polymorphisms in *ftsI* revealed that 131 (18.4%) had a four-AA-insertion between positions 333 and 334 of PBP3, the insertion being particularly prevalent in ST101, ST167, ST410 and ST1284 (Sup. Table 1 and Fig. 5D). Similarly, *ompC* alleles we previously characterized as modifying susceptibility to antibiotics were more prevalent in some STs: an ST38-like *ompC* allele resulting from a recombination event or a G138D mutation were present in 33 and 26 isolates belonging to six different ST, whereas the R195L mutation was observed in 50 isolates all from ST410 (Table S1 and Fig. 5E). In addition, 144 isolates belonging to phylogroups D and F possessed the ST38-like allele inherited vertically. On the other hand the *ompF* gene was pseudogeneized in 47 isolates (Table S1).

To analyze the phylogenetic relationships between F-NRC Cp-*Ec* and other Cp-*Ec* isolates collected worldwide, we built ST-based maximum-likelihood trees for the 14 STs with more than 10 isolates and included to this analysis genome sequences publicly available (Table S2). Together with the analysis of QRDR mutations and mutations in *ftsI* and *ompC*, this showed that 51 (80%) of the ST410 (Fig.S2A), 17 (49%) of the ST167 (Sup Fig.S2B), 3 (33%) of the ST405 (Sup Fig.S2C) and 8 (62%) of the ST101 (Fig. S2D) F-NRC CP-*Ec* belonged to internationally disseminating MDR subclades enriched in CP we previously identified (10). Subclades characterized by mutations in *ftsI* and *ompC,* mainly expressed *bla*_OXA-181_ (ST410) or *bla*_NDM_ (other lineages). Other isolates belonging to these ST were dispersed on their respective phylogenies and with no sign of clonal dissemination for most of them.

ST38 corresponded to the most represented ST (n=92) among the studied CP-*Ec*. The phylogenetic reconstruction together with 314 additional non redundant CP-*Ec* genome sequences retrieved from Enterobase (http://enterobase.warwick.ac.uk/) and 150 non-CP isolates from NCBI provided evidence that 80.4% (n=67) of the French isolates clustered into four clades (Fig. 6). Three of them, G1 (n=35), G2 (n=25) and G4 (n=4) only contained isolates encoding *bla*_OXA-48_, while G3 (n=7), included isolates expressing *bla*_OXA-48_ or its single nucleotide derivative *bla*_OXA-244._ All but one isolates in G1 expressed *bla*_CTX-M-24_, while isolates of G3 expressed *bla*_CTX-M-14b_. The phylogenetic analysis provided evidence of worldwide dissemination of G1, G3 and G4 clades and multiple introduction in France. Strickingly, among the 28 isolates of the G2 clade, 24 were collected in France and four in the Netherlands, suggesting at this time a more regional dissemination. In none of the isolates of the four clades, an IncL plasmid, that generally encode *bla*_OXA-48_ (13), could be identified by PlasmidFinder (14). It suggests a chromosomal integration as previously shown among ST38 isolates collected in the UK (15). Analysis of representative isolates of the G1 and G3 cluster (G1: GCA_005886035.1; G3: GCA_004759025.1), whose genome sequences were completely assembled, confirmed that *bla*_OXA-48_ and *bla*_OXA-244_, in these two isolates respectively, were chromosomally integrated. To assess the genetic support of *bla*_OXA-48_ in the two other clades, we have determined the complete genome sequence of two isolates belonging to G2 (CNR65D6) and G4 (G4:CNR85I8) by combining long-read Pacbio and Illumina sequencing and found that both possess a chromosomally inserted *bla*_OXA-48_ gene. pOXA-48 plasmid sequences of various lengths were cointegrated with *bla*_OXA-48,_ the whole sequence being surrounded by IS*1* insertion sequences with different insertion sites for the four clades (Fig. 7). BLASTN analyses on the assembled Illumina sequences of other isolates of each clade showed that isolates of the same clade share the same insertion sites (see material and methods). A fifth clade, composed of seven closely related French isolates corresponded to a possible outbreak in the East of Paris between December 2014 and December 2015. These isolates are predicted to be highly resistant as they are carrying in addition to *bla*_VIM-4,_ *bla*_CTX-M-15_, as well as 15 to 18 additional ARGs and five mutations in QRDRs.

**Figure 6.**
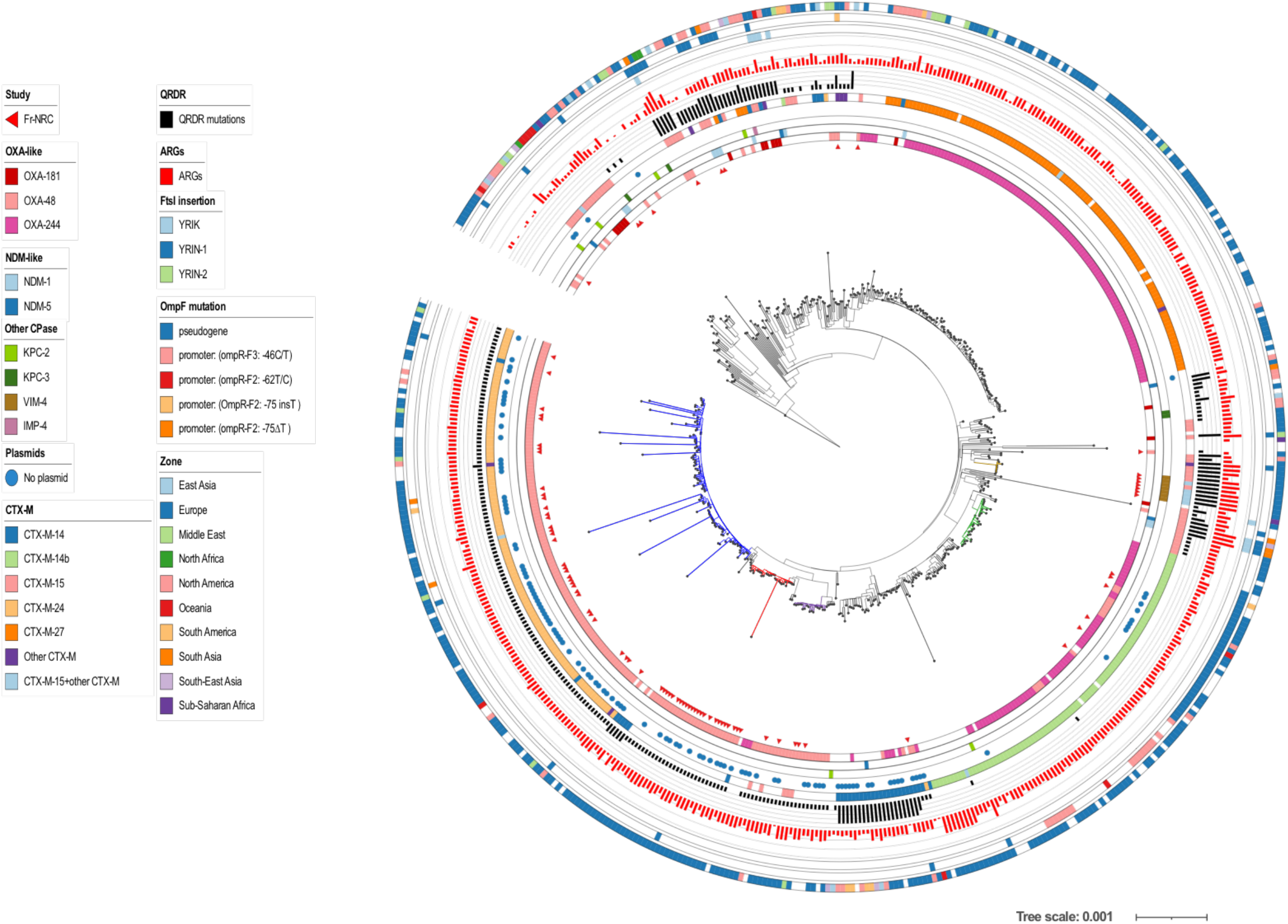
Core genome phylogeny of ST38 isolates. ML Phylogeny was based on genome sequences of 92 CP-*Ec* from the F-NRC and 464 genome sequences retrieved from Enterobase and from the NCBI database including 331 carrying a carbapenemase gene. A core genome (2,900,000 nt) alignment of the *de novo* assemblies on the sequence of GCA_005886035.1 used as a reference was performed by using Parsnp (47); ML phylogeny was built with RAxML (45) from 6,170 core SNPs after removing recombined regions with Gubbins (48). The genome sequence of CNRC6O47 (ST963) was used as an outgroup for the phylogenetic analysis. F-NRC isolates are indicated by red triangles (inner circle). Other genomic features are indicated as indicated in the figure key (left) from the inside to the outside circles: carbapenemases of the OXA, NDM and other types, absence of any plasmid of any Inc-type as identified by using PlasmidFinder (14), ESBL of the CTX-M types, number of mutations in *gyrA* and *parC* QRDRs (FQ resistance), number of ARGs, mutations in *ftsI*, *ompC*, *ompF*, geographical origin. The four OXA-48-like clades (G1, G2, G3, G4) clustering most French isolates are colored in blue, red, violet and green. A fifth clade (G5) corresponding to a possible outbreak of VIM-4 isolates in the East of Paris is colored in brown

**Figure 7.**
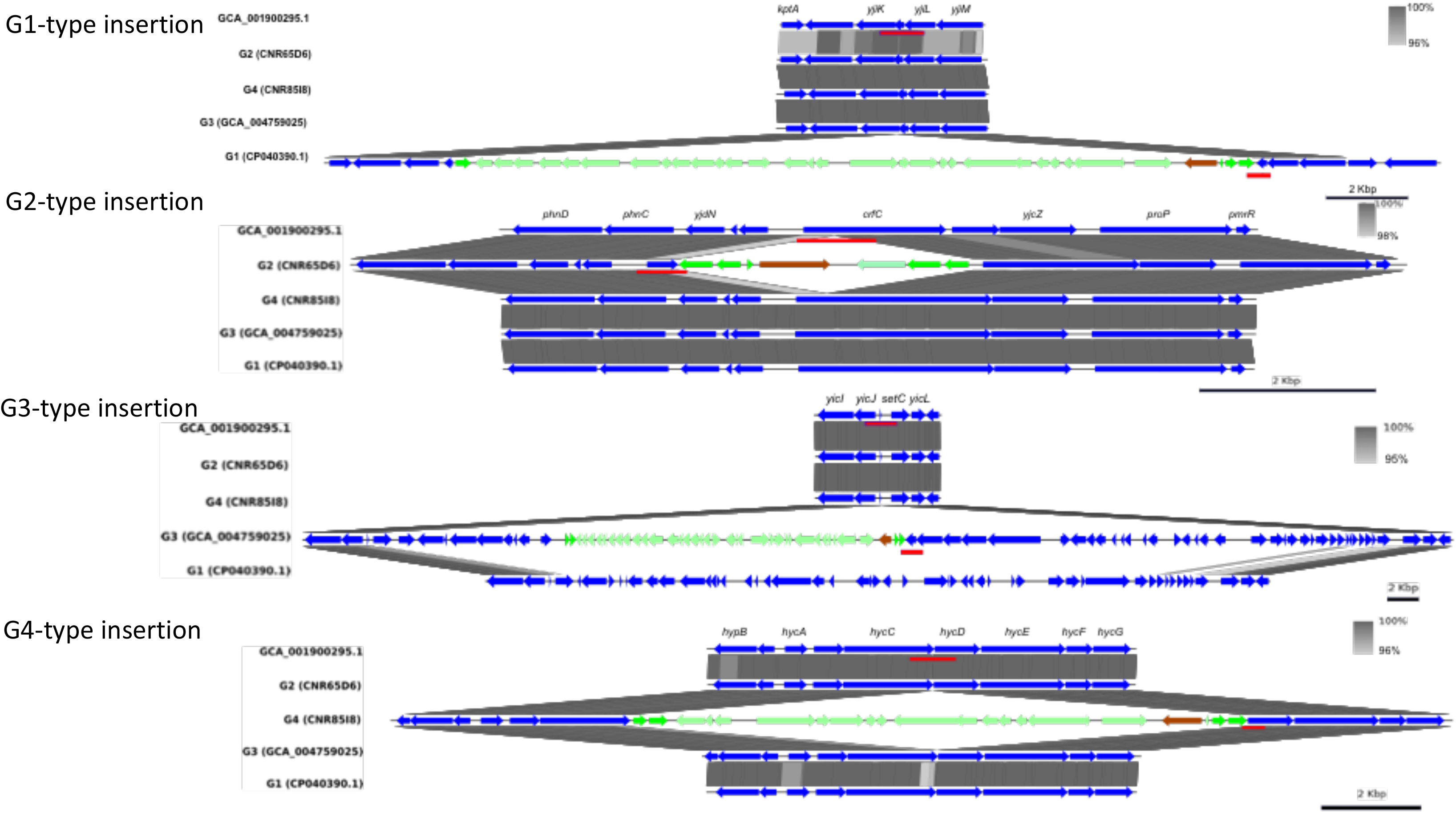
*bla*_OXA-48_ integration sites of ST38 CP-*Ec* clades 1, 2, 3 and 4. The genome sequences of GCA_005886035.1 (CP040390.1), CNR65D6, GCA_004759025.1 (CP038505.1), and CNR85I8 were chosen as representative genomes of clusters 1, 2, 3 and 4, respectively and aligned, by using Blastn, against the sequence of GCA_001900295.1 (CP010116.1), a ST38 isolate that does not carry a carbapenemase gene and whose genome sequence was complete. Gene annotations were derived from MG1655 (NC_000913.3). The regions surrounding the *bla*_OXA-48_ insertion sites in each clade were more precisely aligned against the corresponding sequences of CP010116.1 and of representative genomes of the other clades by using Easyfig (52). *bla*_OXA-48_ is represented by a brown arrow, cointegrated genes of pOXA-48 origin with light green arrows and the IS*1* transposase with a green arrow. Sequences used to perform the BLAST analysis on Illumina contigs are underlined in red. It is to note that, in the G3 clade, *bla*_OXA-48_ integration occurs into a genomic island inserted at tRNASer. Another genomic island is present in G1-type strains but does not carry *bla*_OXA-48_.

ST10 *E. coli* isolates are commensals of a variety of mammals and bird species (16) and studies in different contexts have shown that this ST is also associated with ARGs carriage (17, 18). The 67 F-NRC ST10 CP-*Ec* isolates were distributed throughout the ST10 phylogeny. Nonetheless 31 isolates belonged to a clade enriched with carbapenemase producers (Fig. S3A, in green) that mainly expressed *bla*_OXA-48_, with the exception of three isolates expressing *bla*_OXA-181_, three expressing *bla*_NDM-5_ and one and *bla*_VIM-4_. Two of the *bla*_OXA-181_ and the three *bla*_NDM-5_ expressing isolates were closely related, shared a CTX-M-27 ESBL, four mutations in QRDR and a *ftsI* allele with a four AA YRIN insertion (Fig. S3A, in red).

ST131 isolates are major contributors for extra-intestinal infections and the main responsible for the dissemination of CTX-M-15 ESBL (19, 20). ST131 evolution has been thoroughly analyzed and four main lineages A, B, C1 and C2 have been described (21). We identified only 17 ST131 CP-*Ec* isolates received by the F-NRC (Fig. 2), ten belonged to lineage A, while the others were scattered in the three other lineages (Fig. S3B). The main carbapenemase associated with these isolates was OXA-48 (N=10). We also did not observe any association with the carriage of *bla*_CTX-M_ genes, as only four ST131 CP-*Ec* isolates carryied a *bla*_CTX-M-15_ gene. These results are in accordance with the UK survey (8) and our previous observations from sequences retrieved from public databases (10).

ST940 (phylogroup B1) and ST1284 (phylogroup A) represented 2.4% (n=17) and 1.9% (n=14) of the sequenced isolates while they were poorly represented in sequence databases. For instance, in May, 2021, only 83 and 103 sequences genome sequences could be retrieved from Enterobase for these ST respectively, but with 43% (n=36) and 22% (n=22) carrying a carbapenemase gene respectively. This suggested that ST940 and ST1284 were associated with the dissemination of carbapenemase genes. Phylogenetic trees were drawn by adding the non-redundant sequences from enterobase to those of the F-NRC and NCBI. For ST940 (Fig. S4A) this revealed a well-differentiated sub-clade of 18 isolates from Asia, Australia and Europe carrrying *bla*_NDM-5_ and sharing mutations in *ftsI* and *ompC*. Isolates from the F-NRC were distributed on the tree, showing that the over-representation of ST940 in France did not result from local outbreaks. Three F-NRC isolates belonged to a second smaller sub-clade of *bla*_NDM_ expressing isolates, also mutated in *ftsI* and o*mpC*. In contrast, in ST1284, thirteen encoding *bla*_OXA-181_, were found to be closely related (Fig. S4B). Metadata analysis showed that 12 out of them were recovered from the same health facility in the Paris suburb during a one-month period in 2015, demonstrating a local outbreak origin. A fourteenth unrelated isolate expressed *bla*_NDM-5_. The F-NRC isolates belonged to a clade characterized by four mutations in QRDR determinants, a YRIN duplication in PBP3, a G137D mutation in OmpC and containing additional OXA-181- or NDM-producing isolates. Among the fifteen ST90 CP-*Ec* isolates, nine encoding the *bla*_OXA-204_ allele were isolated in hospitals East of Paris between August 2012 to April 2013 and suspected to be associated with the use of a contaminated endoscope (22). Isolates closely related to this outbreak reappeared in 2014 (three times), and in 2015 (twice), also in East of Paris located hospitals. They belonged to a MDR clade characterized by four mutations in QRDR (Fig. S5A). Finally, in the four last STs with more than 10 CP-*Ec* isolates (ST69, ST88, ST617, and ST648), French CP-*Ec* isolates were distributed throughout their respective phylogenetic trees (FigS5B and C, S6A and B) with no sign of clonal dissemination, except for a small cluster of five ST69 from two geographical regions and expressing *bla*_OXA-48_.

In addition to potential outbreaks detected in ST38 (*bla*_VIM-4_), in ST90 (*bla*_OXA-204_) and ST1284 (*bla*_OXA-181_), we also found evidence for another potential outbreak among isolates belonging to the ST359. The five isolates encoded *bla*_OXA-48_ and *bla*_CTX-M-32_ and had three mutations in QRDR. They were isolated during January 2014, four of them in the south-eastern part of France and one in the Parisian suburbs.

## DISCUSSION

The prevalence of CP-*Ec* is increasing worldwide. However whether this reveals conjugation event in *E. coli* isolates by circulating CP-encoding plasmids or the emergence and dissemination of at-risk clones is still largely unknown. Here, we have analyzed, by using WGS, 691 (87.8%) of the CP-*Ec* isolates received by the F-NRC during the period 2012-2015 and 22 isolates from the Bicêtre hospital strain collection (2001-2011) to characterize both the diversity of carbapenemase genes and of CP-*Ec* isolates in France. Altogether 713 CP-*Ec* were sequenced, representing to our knowledge the most extended collection of CP-*Ec* sequenced, published to date.

The number of CP-*Ec* isolates sent to the F-NRC was found to strongly increase during the four-year period of analysis, which might be a consequence of an increased circulation of CP-*Ec* in France but also of an increased screening of potential CPE carriers at their admission at hospital. Indeed, while the number of infection-related CP-*Ec* isolates was regularly increasing, their proportion compared to the total number of received and sequenced isolates decreased by two-fold, with a clear change observed between 2013 and 2014 (Fig. 1). This is likely a consequence of the implementation of the recommendations on MDR screening of the French Public Health Advisory Board 2013 (23). However, as CP-*Ec* isolates were sent on a voluntary basis by clinical laboratories, we cannot exclude some sampling bias.

The most predominant carbapenemase allele was *bla*_OXA-48_ (65%) followed by *bla*_OXA-181_ (14.1%) which detection constantly increased from no cases in 2012 to 73 in 2015. Next was *bla*_NDM-5_ that was found to progressively substitute to the *bla*_NDM-1_ gene in frequency of detection (Fig. 4). The predominance of *bla*_OXA-48-like_ genes and particularly of *bla*_OXA-48_, among CP-*Ec* was also noted in other studies (9, 24,25) although in lower proportions. The small number of *bla*_KPC_ genes detected in these studies is in contrast with the situation reported in *K. pneumoniae* for many European countries (26). However, *bla*_OXA-48_ was found the most prevalent allele in *K. pneumoniae* in France (2) and in the Netherlands (9). This difference might result from a lower capacity for *bla*_KPC_ encoding plasmids to conjugate to *E. coli* as compared to pOXA-48 or to a higher fitness cost. It might also be linked to sampling differences as different thresholds of carbapenem susceptibility might have been used to collect CRE isolates among studies. Indeed OXA-48-like carbapenemases are generally associated with lower levels of carbapenem resistance than KPC. This analysis was performed on isolates collected between 2012 and 2015 and revealed early trends in the evolution of CP-*Ec* isolated in France. It will be essential in the characterization of the genomic evolution of CP-*Ec* in more recent years, which remains to be performed.

The number of ARGs identified in the CP-*Ec* isolates from the F-NRC varied from one (the carbapenemase gene only) to 26, showing that carbapenemase genes were acquired not only in MDR isolates but also in an *E. coli* population that can be considered as "naïve" relative to an evolution under antibiotic selective pressure and resident of the intestinal microbiota. This was particularly true for isolates producing *bla*_OXA-48_ that were generally associated with a lower number of resistance genes than isolates producing other carbapenemases, irrespectively of their clinical status (Fig. 4A). In particular 118 out of the 121 (97.5%) isolates with no other ARG than the carbapenemase gene carried *bla*_OXA-48_. Therefore, the predominance of *bla*_OXA-48_ among the carbapenemase genes identified in the F-NRC collection might mainly rely on the higher conjugative transfer rate of pOXA-48 related plasmids compared to those encoding other carbapenemases (27). pOXA-48 has indeed been shown to rapidly conjugate among *Enterobacterales* in hospitalized patients (28). This could also explain the high frequency of *bla*_OXA-48_ carrying isolates belonging to ST10, ST69, ST88, ST127, ST12 and ST58 (Fig. 5), that carried a small number of resistance genes and QRDR mutations and among which four were previously identified among the most frequent *E. coli* ST characterized from fecal samples (29).

On the other hand, MDR CP-*Ec* isolates may result from the transfer of carbapenem resistance genes into isolates already selected through multiple exposures to antibiotic treatments that have already acquired multi-resistance plasmids or chromosomal mutations increasing their intrinsic drug resistance, such as mutations in QRDR, in *ftsI* and/or in porin genes. Alternatively, the circulation of carbapenem-resistant lineages, such as the OXA-181-producing ST410 *E. coli* lineage, showing a stable association between the lineage and the resistance gene, may also occur and significantly contribute to the number of CP-*Ec* isolates collected. Discriminating between both alternatives would require a more extended comparison of plasmids carried in these lineages after long-read sequencing similarly to what been done in *K. pneumoniae* (30). Strikingly, the analysis of *ftsI* alleles coding for a PBP3 with the four amino-acid duplication revealed a contrasted situation among CP-*Ec* isolates collected in France, with the duplication in *ftsI* found in 72.3%, 77.8% and 67% of the isolates carrying *bla*_NDM-5_ (34/47), *bla*_NDM-7_ (7/9) and *bla*_OXA-181_ (68/101) genes but in only 0.7% (4/458) of the isolates carrying *bla*_OXA-48_. We previously proposed that the fixation of mutations reducing the intrinsic susceptibiliy to carbapenems might have favored the efficient conjugative transfer of plasmids carrying the carbapenemase genes from other CPE species in the gut (10). Transconjugant would be selected in a context of low biliary excretion of carbapenems or other ß-lactams after parenteral administration. Increasing the proportion of the donor and recipient bacteria would therefore be less necessary for plasmids with high conjugative rates, such as pOXA-48 (27). Alternatively, these mutations might also have favored the plasmid maintenance by increasing the resistance level and selection of the isolates during ß-lactam or carbapenem treament. Of particular concern in France are the OXA-181 producing ST410 and the NDM (NDM-1, −5 and −7) ST167 lineages that significantly contribute to the circulation of carbapenem resistance. However our study also reveals smaller lineages, that although less frequently encountered in international studies, are nevertheless circulating. In contrast we did not obtain evidence for clonal dissemination of CP-*Ec* ST131 lineages derived from the B2 clade responsible for *bla*_CTX-M-15_ gene dissemination (19, 20).

Despite their high frequency among the F-NRC collection, the ST38 isolates do not enter into one of the two previous categories. Indeed, while they mainly express *bla*_OXA-48_ or its single-point-mutant derivative *bla*_OXA-244_ (31), our phylogenetic analysis revealed that a majority of them belong to four different lineages, including one mostly associated with a rapid dissemination in France. In contrast with OXA-181 and NDM-producing lineages previously described, these lineages are associated with a moderate number of ARGs and QRDR mutations. None of the ST38 isolates is mutated in *ftsI*, however we previously demonstrated that all are characterized by an *ompC* allele that encodes a porin with a reduced permeability to certain ß-lactams and has disseminated into unrelated lineages by homologous recombination (10). A common feature of the four lineages is the integration of part of the pOXA-48 plasmid carrying *bla*_OXA-48_. This could have reduced a fitness cost of this plasmid and facilitated the clonal expansion of these lineages. However, ST38 isolates were identified as unfrequent colonizer of the GI tract (29, 32). Therefore additional features of these CP-*Ec* ST38 lineages might have contributed to their dissemination.

In conclusion, by analysing genome sequences of CP-*Ec* isolates collected by the F-NRC we showed that MDR lineages, enriched in carbapenemase-producing isolates are circulating in France, some of them, such as ST1284 isolates being associated with outbreaks. It also suggests that the evolutionary trajectory may depend on the carbapenemase gene, *bla*_OXA-181_ or *bla*_NDM_ genes being more frequently associated with the evolution of MDR *E. coli* lineages characterized by mutations in *ftsI* and *ompC*. Surveillance of these mutations may be an important parameter in controling the circulation of MDR lineages. On the other hand, carbapenemase genes are also frequently acquired through plasmid dissemination from other bacterial species. Depending on the resistance background of the receiver *E. coli*, this may lead to XDR or to isolates sensitive to a broad range of antibiotics. Finally, we also observed a strong and rapid dissemination of ST38 isolates that might have been favored by a reduced susceptibility to carbapenems linked to the ST38 *ompC* allele and by the chromosomal integration of the carbapenemase gene. These results strengthen our model of different evolutionary trajectories associated with the gain of carbapenemase genes (10). They also show that systematic genome sequencing of CP-*Ec* and at a larger of CPE isolates, irrespective of their clinical or resistance status is able to provide useful information not only on the circulation of MDR lineages, but also on the propagation of resistance genes through horizontal gene transfer.

## MATERIAL AND METHODS

### Isolate collection and sequencing

CP-*Ec* isolates analyzed in this study were collected by the F-NRC, mainly between the years 2012 to 2015. Twenty-two additional isolates we have received before the creation of the F-NRC in 2012, were included in the analysis. Information on these isolates as the year of isolation, the region and department in France and summary of their genomic features are reported in Table S1.

### Whole genome sequencing and analyses

DNA were extracted by using the Qiagen Blood and Tissue DNA easy kit. Sequencing libraries were constructed by using the Nextera XT kit (Illumina) following the manufacturer instruction, and sequenced with the Illumina HiSeq2500 or NextSeq500. FastQ files were trimmed for adaptors and low-quality bases (setting the minimum base quality threshold to 25) with the Cutadapt fork Atropos (33). *De novo* assemblies were generated from the trimmed reads with SPAdes v3.12.0 (34), using k-mer sizes of 51, 71, 81, and 91, the coverage cut-off option was set to “auto” and the “careful” option was activated. QUAST v2.2 (35) was used to assess the assembly quality and contigs shorter than 500 bp were filtered out for phylogenetic analyses. The ST38 isolates CNR65D6 and CNR85I8 were sequenced to completion by using the long-read Pacbio technology. PacBio reads were assembled with the RS_HGAP_Assembly.3 protocol from the SMRT analysis toolkit v2.3 (36) and with Canu (37), polished with Quiver (36) and manually corrected by mapping Illumina reads using Breseq (38). Assembled genomes were annotated with Prokka v1.9 (39). To analyze the F-NRC CP*Ec* isolates belonging to the main ST recovered during the analysis in a more global context, their genome sequences were combined to the assembled genome sequences from the same ST retrieved from the NCBI database (July, 2019) and in some STs from Enterobase (http://enterobase.warwick.ac.uk/, April 2021). Retrieved genomes were annotated in the same way as the genome sequences of the F-NRC isolates (Table S2).

To generate a core-genome phylogeny tree of all isolates, the best reference genome was first selected among a set of 18 genomes analyzed by Touchon et al. (40) using the software refRank (41) and three subsets of 100 randomly selected sequences from our study as an input. This led to the selection of the genome sequence of strain MG1655-K12 (NC_000913.3) as the reference for read mapping and SNP identification. Sequence reads for the 17 reference genomes (40) as well as for the genome sequence of *Escherichia fergunsonii* strain ATCC 35469 (NC_011740.1), used as an outgroup were simulated with ART (42). Trimmed sequencing reads were mapped against the MG1655-K12 genome with BWA-MEM algorithm of the BWA v0.7.4 package (43). For SNP calling the Genome Analysis Toolkit (GATK) v3.6.0 (44) was used with the following criteria, a minimum depth coverage (DP) of 10, a quality by depth (QD) bigger than 2, a fisher strain bias (FS) below 60, a root mean square of the mapping quality (MQ) above 40, and the mapping quality rank sum test (MQRankSum) and the read position rank sum test (ReadPosRankSum) greater than −12.5 and −8 respectively. A Maximum-Likelihood tree was estimated with RAxML v8.28 (45) using core-genome SNPs after removing positions in the accessory genome identified with the filter_BSR_variome.py script from the LS-BSR pipeline (46). For ST with more than 10 CP-*Ec* isolates from the F-NRC, a core genome alignment was generated with Parsnp (47), by using a finished genome sequence as reference. A closely related isolate outside the ST lineage was selected from the global phylogenetic tree including all F-NRC isolates and used as outgroup to root the ST phylogenetic trees. Maximum-Likelihood (ML) trees were generated with RAxML v8.28 (45) after removing regions of recombination with Gubbins (48). All graphic representations were performed by using ITOL (49).

### MLST type, phylogroups, resistance gene and plasmid replicon identification

Sequence type was assigned to each assembly through a python script that relies on BLAST (10). Antibiotic resistance genes (ARGs) and mutations in *gyrA, parC* and *parE* quinolone resistance determining regions (QRDR) were identified with Resfinder 4.0 and PointFinder (50) run in local respectively. The scripts and database (Retrieved: January 4, 2021) were downloaded from the repositories of the Centre for Genomic Epidemiology (https://bitbucket.org/genomicepidemiology/). The identified ARGs were manually reviewed to eliminate potential redundant ARG predicted at the same genomic position. *mdf*(A) that is present in *E. coli* core genome was not taken into account. Phylogroups were assigned by using EzClermont (51) run in local. Plasmid replicons were identified with plasmidfinder run on each assembly (14). *ftsI*, *ompC and ompF* CDS sequences and *ompF* promoter sequences were identified by using BlastN. Translated or nucleotide sequences were clustered by cd-hit (cd-hit-v4.8.1) with an amino acid (FtsI, OmpC, OmpF) or nucleotide sequence (*ompF* promoter) identity threshold of 1. Sequences of each cluster were aligned to detect mutations in regions of interest: four amino-acid insertions between P333 and Y334 and E349K and I532L mutations (FtsI), mutation modifying charge in L3 constriction loop, R195L mutation, nonsense or frameshift mutations or OmpC sequence clustering with ST-38 OmpC sequences (OmpC), nonsense mutations (OmpF) or mutations affecting one of the OmpR-boxes: mutation −46T/C in OmpR-F3 box and mutation −75ΔT in OmpR-F2 box (*ompF* promoter).

### Determination of *bla*_OXA-48_ insertion site in ST38 clades 1, 2, 3 and 4

The genome sequences of GCA_005886035.1 (CP040390.1), CNR65D6, GCA_004759025.1 (CP038505.1), and CNR85I8 were chosen as representative genomes of clusters 1, 3, 2 and 4, respectively and aligned, by using Blastn, against the sequence of GCA_001900295.1 (CP010116.1), a ST38 isolate that does not express a carbapenemase and whose genome sequence was complete. The regions surrounding the *bla*_OXA-48_ insertion sites in each clade were more precisely aligned against the corresponding sequences of other representative genomes by using Easyfig (Easyfig_mac_2.1) (52) (Fig. S3). To perform a blast analysis of insertion sites by using contigs generated from Illumina sequences, smaller sequences were chosen as queries as following: nucleotides (nt) 1240133..1240934, nt 964173..965072, nt 419599..421606, nt 4206783..4207690 from GCA_001900295.1 to screen for native sequences corresponding to the four *bla*_OXA-48_ insertion site in G1, G2, G3 and G4 clades and sequences nt 4460332..4460759 from CP040390.1, nt 4763987..4764506 from CNR65D6, nt 100784..102087 from CP038505.1 and nt 1206426..1206985 from CNR85I8 to screen for the integration site plus IS*1* insertion as occurring in G1, G2, G3 and G4 clades respectively.

### Statistical analysis

Pearson’s chi-squared tests were performed by using standard libraries contained within the R statistics package (http://www.R-project.org)

### Availability of data

All sequence data have been deposited at DDBJ/EMBL/GenBank (BioProject PRJEB46636) and bioSample identifiers for the Illumina sequence data are listed in Table S1. Complete genome sequences of CNR65D6 and CNR85I8 and the long-read sequencing data have been deposited at DDBJ/EMBL/GenBank with the accession number ERZ3517884 (PacBio reads, ERR6414227) and ERZ3518335 (ERS6682837 PacBio reads) respectively.

## ACKNOWLEDGMENTS

This work was supported by grants from the French National Research Agency (ANR-10-LABX-62-IBEID, ANR-10-LABX-33, INCEPTION project (PIA/ANR-16-CONV-0005)) and from the European Union’s Horizon 2020 research and Innovation Program under grant agrement No 773830 (Project ARDIG, One Health EJP). Pengdbamba Dieudonné Zongo is a scholar of Ed525 CDV, Sorbonne Université, Paris.

---

**Figure S1.**
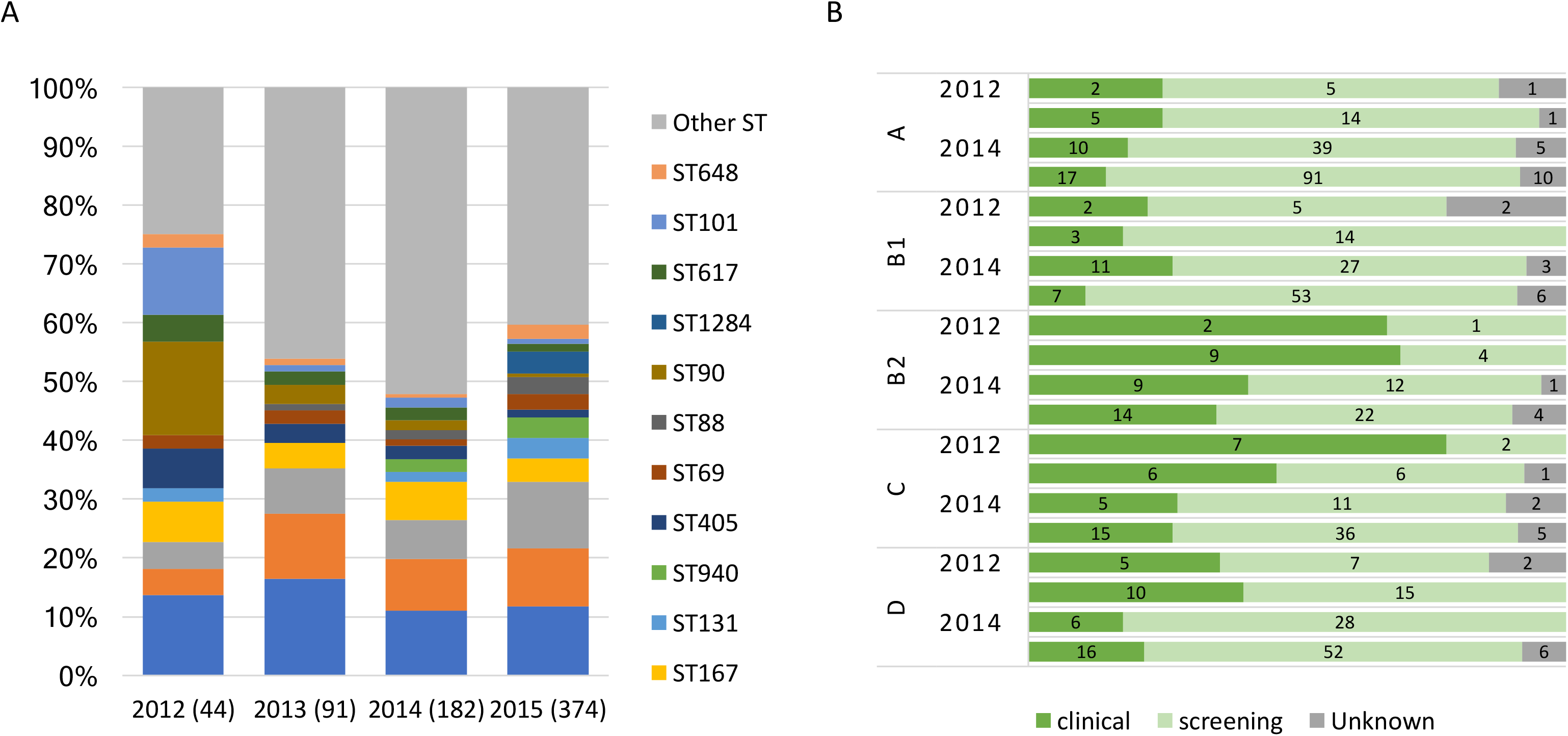
Per-year analysis of the origin of the isolates received by the F-NRC. **A**. Proportions of the isolates belonging to the different ST. The total number of isolates is indicated between brackets. The large proportion of ST90 isolates collected in 2012 is linked to a local outbreak. **B**. Proportions of the isolates collected in screening and infection situations as a function of the year and the phylogroup. The absolute number of isolates for each class is indicated inside the bars.

**Figure S2.**
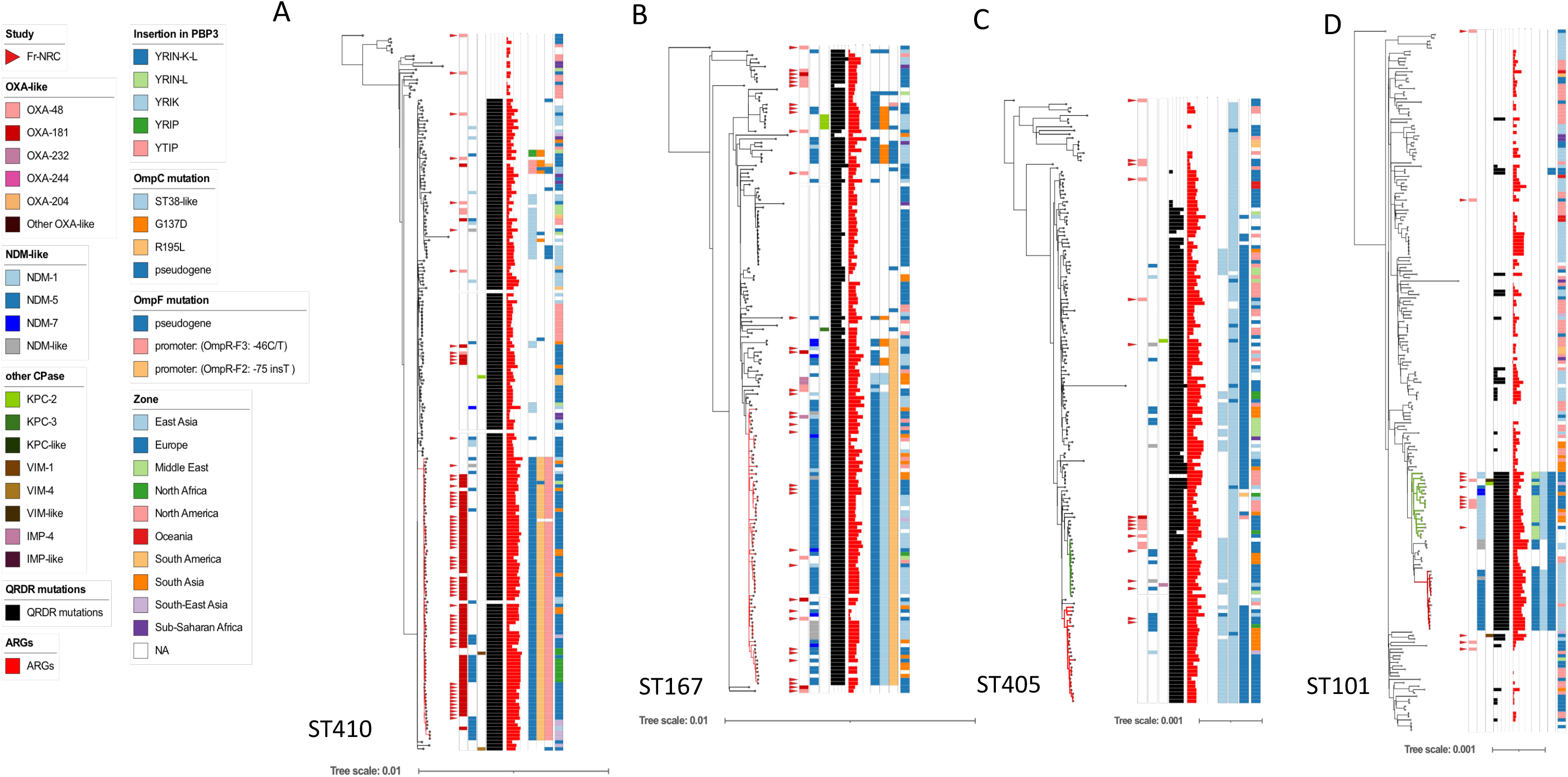
Core genome phylogenies of the main ST characterized by clades disseminating internationally. A. ST410; B. ST167; C. ST405; D. ST101. Phylogenies were based on genome sequences of A: 64 CP-*Ec* from the F-NRC and 146 genome sequences from the NCBI database;B: 35 CP-*Ec* from the F-NRC and 134 genome sequences from the NCBI database; C. 15 CP-*Ec* from the F-NRC and 145 genome sequences from the NCBI database; D. 13 CP-*Ec* from the F-NRC and 194 genome sequences from the NCBI database. Core genome (ST410: 3,522,000 nt; ST167: 3,276,000 nt; ST405: 3,685,000 nt; ST101: 3,774,000 nt) alignments of the *de novo* assemblies on the sequences of GCA_001442495.1 (ST410), GCA_003028815.1 (ST167), GCA_002142675.1 (ST405), GCA_002163655.1 (ST101) used as reference were performed by using Parsnp; ML phylogeny was built with RAxML from 6,176 (ST410), 10,393 (ST167), 37,482 (ST405), 19,967(ST101) core SNPs after removing recombined regions with Gubbins. The genome sequences of CNR93E7 (ST88), CNR93D10 (ST746), CNR73I9 (ST115), CNR93I2 (ST906) were used as outgroups for the phylogenetic analyses of ST410, ST167, ST405 and ST101 respectively. The origin from the F-NRC is indicated by red triangles close to the isolate name. Other genomic features are indicated as indicated in the figure key (left) from the inside to the outside lines: carbapenemases of the OXA, NDM and other types, number of mutations in *gyrA* and *parC* QRDR region (FQ resistance), number of ARGs, mutations in *ftsI*, *ompC*, *ompF*, geographical origin.

**Figure S3.**
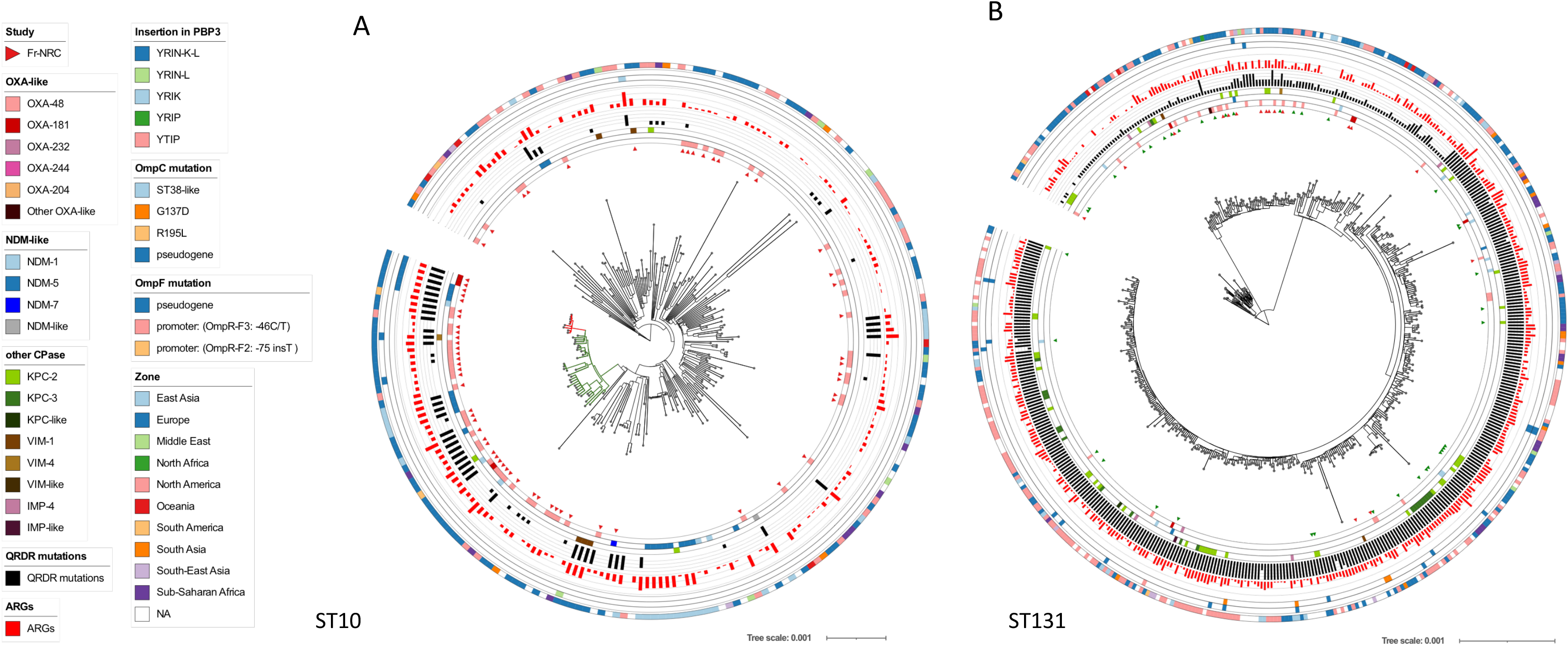
Core genome phylogenies of ST10 and ST131 isolates. **A.** ST10 phylogeny based on the genome sequences of 67 CP-*Ec* from the F-NRC and 153 genome sequences from the NCBI. **B.** ST131 phylogeny based on the genome sequences of 17 CP-*Ec* from the F-NRC and 462 genome sequences from the NCBI. Core genome (ST10: 813,000 nt; ST131: 1,641,000 nt) alignments of the *de novo* assemblies on the sequence of MG1655 (ST10) or GCA_000285655.3 (ST131) used as references were performed by using Parsnp; ML phylogeny was built with RAxML from 20,245 (ST10) and 14,366 (ST131) core SNPs after removing recombined regions with Gubbins. The genome sequences of CNR93D10 (ST746) and CNRAL47G10 (ST640) were used as outgroups for the phylogenetic analyses of ST10 and ST131, respectively. F-NRC isolates are indicated by red triangles (inner circle). Other genomic features are indicated as indicated in the figure key (left) from the inside to the outside circles: carbapenemases of the OXA, NDM and other types, number of mutations in *gyrA* and *parC* QRDR region (FQ resistance), number of ARGs, mutations in *ftsI*, *ompC*, *ompF*, geographical origin. The ST10 clade enriched in CP-*Ec* is indicated in green and the sub-clade with the YRIN insertion in PBP3 in red. Note that the circle for *ftsI* mutations is absent in ST131, as no YRIN-like duplication in PBP3 was identified among the analyzed isolates.

**Figure S4.**
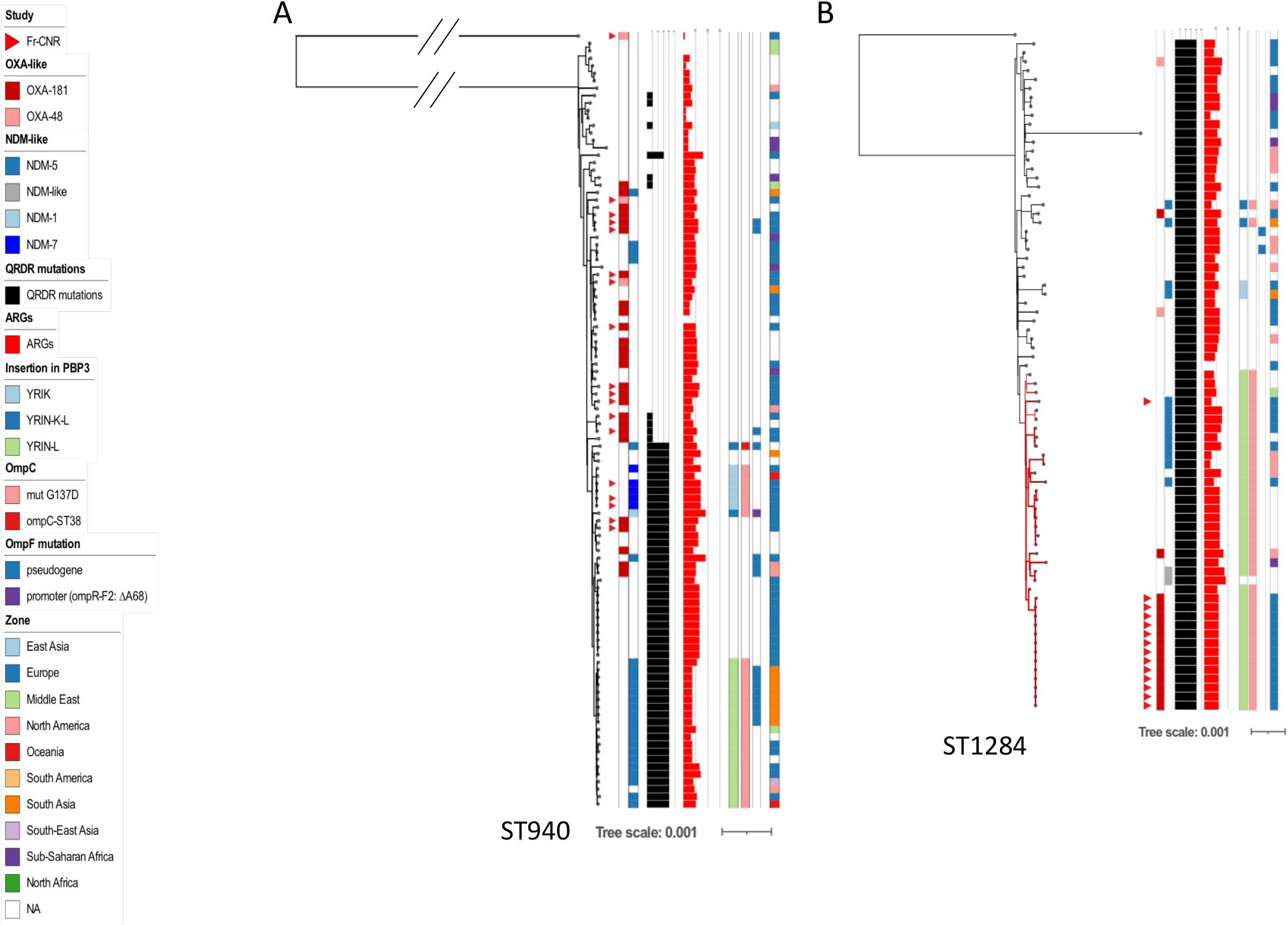
Core genome phylogeny of ST940 and ST1284 isolates. **A:** ST940 phylogeny based on genome sequences of 17 CP-*Ec* from the F-NRC and 88 genome sequences from Enterobase and the NCBIdatabase **B:** ST1284 phylogeny based on genome sequences of 14 CP-*Ec* from the F-NRC and 60 genome sequences from Enterobase and the NCBI database. A core genome (ST940: 3,639,000 nt; ST1284: 3,779,000 nt) alignment of the *de novo* assemblies on the sequence of ST940: ESC_LB2149 and ST1284: ESC_LB2152AA (from Enterobase) used as references was performed by using Parsnp; ML phylogeny was built with RAxML from 13,859 (ST940) and 2,009 (ST1284) core SNPs after removing recombined regions with Gubbins. The genome sequences of CNR98G1 (ST3022) and MG1655 (ST10) were used as outgroups for the phylogenetic analyses of ST940 and ST1284, respectively. F-NRC isolates are indicated by red triangles first column on the left. Other genomic features are as indicated in the figure key (left) from the left to the right columns: carbapenemases of the OXA, NDM and other types, number of mutations in *gyrA* and *parC* QRDR (FQ resistance), number of ARGs, mutations in *ftsI*, *ompC*, *ompF*, geographical origin.

**Figure S5.**
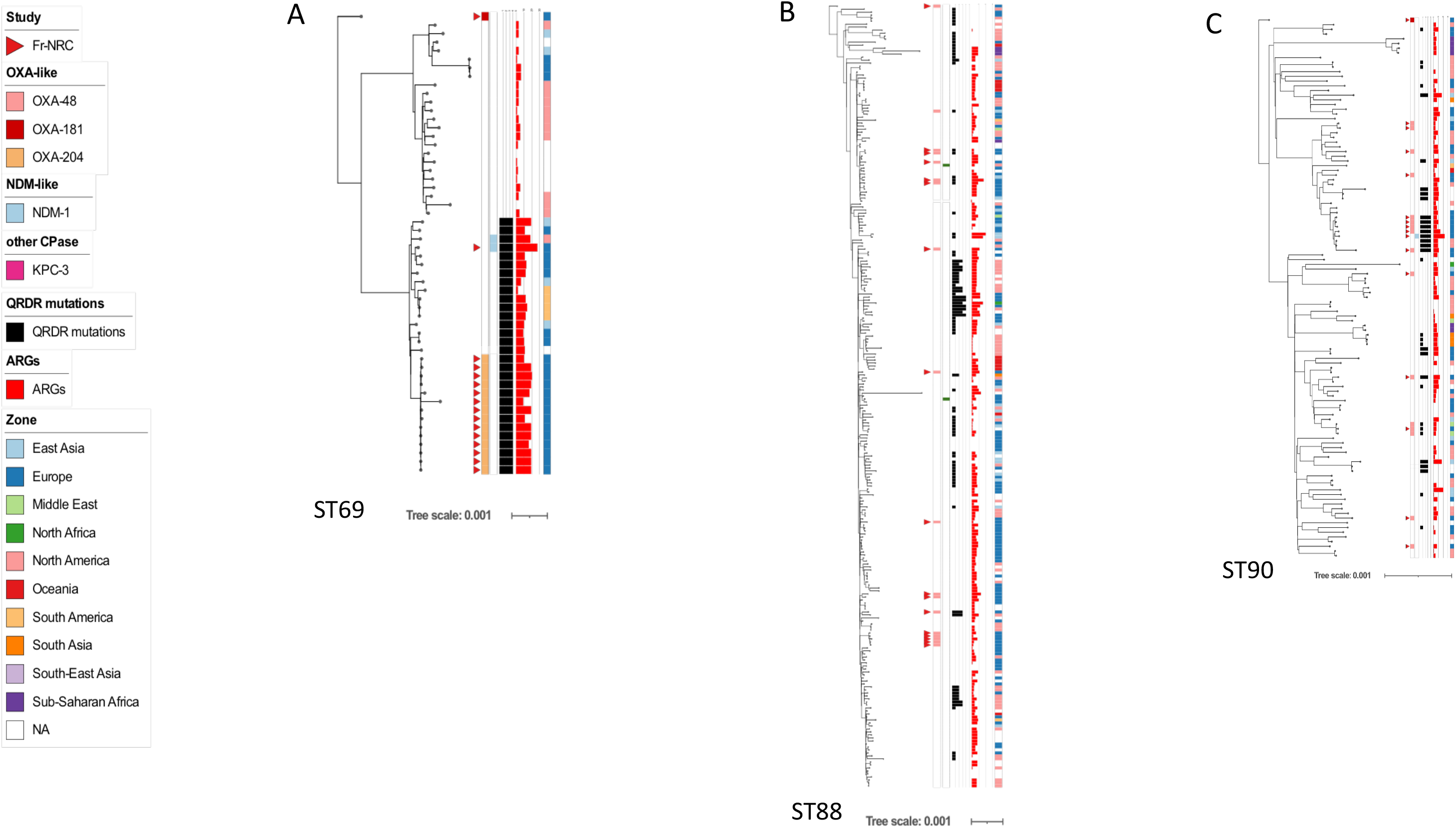
Core genome phylogenies of ST69, ST88 and ST90. isolates. **A:** ST90 phylogeny based on the 15 CP-*Ec* from the F-NRC and 38 genome sequences from the NCBI database**; B:** ST69 phylogeny based on the 16 CP-*Ec* from the F-NRC and 244 genome sequences from the NCBI database; **C.** ST88 phylogeny based on the 15 CP-*Ec* from the F-NRC and 99 genome sequences from the NCBI database. Core genome (ST69: 2,735,000 nt; ST88: 3,472,000 nt; ST90: 3,801,000nt) alignments of the *de novo* assemblies on the sequence of GCA_002443135.1 (ST69), GCA_002812685.1 (ST88) or GCA_001900635.1 (ST90) used as references were performed by using Parsnp; ML phylogeny was built with RAxML from 4,596 (ST69), 12,557 (ST88) or 2,878 (ST90) core SNPs after removing recombined regions with Gubbins. The genome sequences of CNR33D9 (ST394), CNR83B9 (ST410) and CNR88B9 (ST847) were used as outgroups for the phylogenetic analyses of ST69, ST88 and ST90, respectively. F-NRC isolates are indicated by red triangles (first column on the left). Other genomic features are indicated as indicated in the figure key (left) from the left to the right columns: carbapenemases of the OXA, NDM and other types, number of mutations in *gyrA* and *parC* QRDR (FQ resistance), number of ARGs, mutations in *ftsI*, *ompC*, *ompF*, geographical origin.

**Figure S6.**
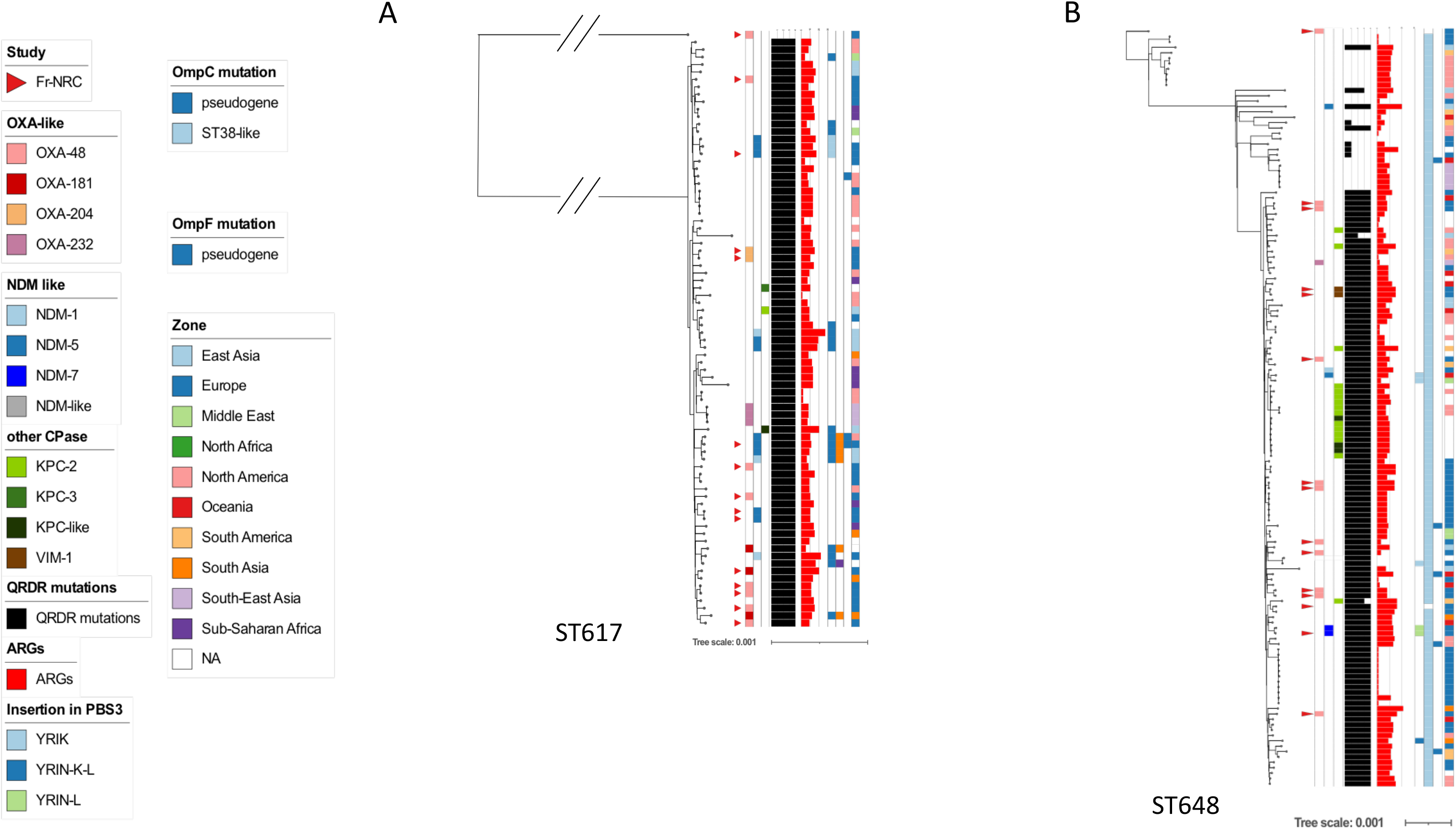
Core genome phylogeny of ST617 and ST648 isolates. **A:** ST617 genome sequences of 15 CP-*Ec* from the F-NRC and 65 genome sequences from the NCBI database; **B:** ST648 genome sequences of 14 CP-*Ec* from the F-NRC and 126 genome sequences from the NCBI database. A core genome (ST617: 3,407,000; ST648: 3,501,000 nt) alignment of the *de novo* assemblies on the sequences of GCA_002142695.1 (ST617) or GCA_004138645.1 (ST648) used as references was performed by using Parsnp; ML phylogeny was built with RAxML from 15,030 (ST617) or 3,782 (ST648) core SNPs after removing recombined regions with Gubbins. The genome sequences of CNR93D10 (ST746) and CNR71A8 (ST1485) were used as outgroups for the phylogenetic analyses of ST617 and ST648, respectively. F-NRC isolates are indicated by red triangles close to the isolate name. Other genomic features are indicated as indicated in the figure key (left) from the left to the right columns: carbapenemases of the OXA, NDM and other types, number of mutations in *gyrA* and *parC* QRDR (FQ resistance), number of ARGs, mutations in *ftsI*, *ompC*, *ompF*, geographical origin.

## Notes

### Competing Interest Statement

The authors have declared no competing interest.

### Summary of Updates

This version of the manuscript has been revised with an additional figure showing the four insertion sites of pOXA-48 in the 4 ST38 sub-lineages carrying blaOXA-48

